# Caught in transition: facultative intracellularity and genome evolution of *Symbiopectobacterium* in *Rhodnius* species

**DOI:** 10.64898/2026.08.16.744597

**Authors:** Tanisha Moons, Sandra Yadira Mendiola, Hassan Tarabai, Vaclav Hypsa, Kevin J. Vogel, Eva Novakova

## Abstract

Blood-feeding insects typically depend on obligate intracellular bacterial symbionts that provide essential B vitamins absent from vertebrate blood. In contrast, kissing bugs (Triatominae) have long been considered atypical because they rely primarily on extracellular gut-associated bacteria. Recent reports of the genus *Symbiopectobacterium* in *Rhodnius* species raise questions about the diversity and evolution of symbiosis in these insects. Here, we investigate the distribution, genome evolution, and tissue localization of *Symbiopectobacterium* in the genus *Rhodnius*. Using comparative metagenomics, phylogenomics, fluorescence *in situ* hybridization, transmission electron microscopy, and hemolymph screening, we characterize a *Symbiopectobacterium* genome from *Rhodnius prolixus* and assess its occurrence across publicly available datasets representing multiple *Rhodnius* species. The *R. prolixus* strain possesses a large, highly dynamic genome enriched in mobile genetic elements, pseudogenes, and remnants of secretion systems, while retaining biosynthetic pathways for several B vitamins. Comparative analysis revealed variation in genome reduction among *Rhodnius*-associated strains, suggesting ongoing and potentially independent transitions toward host-restricted symbiosis. Localization analyses detected *Symbiopectobacterium* intracellularly within posterior midgut epithelial cells and occasionally in the hemolymph, consistent with a facultative intracellular lifestyle. However, no bacteriomes or stable intracellular structures were observed. Together, these findings indicate that *Symbiopectobacterium* represents an intermediate stage in the transition from environmentally associated bacteria to obligate intracellular mutualists in Triatominae.

## Introduction

Obligate blood-feeding insects, which feed exclusively on vertebrate blood throughout all developmental stages, typically harbor obligate bacterial symbionts (Hosokawa et al., 2010; Hosokawa & Fukatsu, 2020; Lalzar et al., 2014; Sonenshine & Stewart, 2021; Zhong et al., 2007), whose role is to compensate for the low B-vitamin content of vertebrate blood (Baines, 1956; Duncan, 1926; Duron & Gottlieb, 2020). These symbionts share common features across diverse host lineages, reflecting their evolutionary transition from free-living bacteria to obligate mutualists. Hallmarks of this lifestyle include strict vertical transmission and host– symbiont coevolution, intracellular residence within specialized host cells or organs (bacteriomes), and the provision of essential metabolic services to the host, such as the synthesis of nutrients unavailable from the host diet. At the genomic level, this evolution is marked by extreme genome reduction, AT enrichment, and loss of redundant metabolic capacities (Douglas, 2015; Moran, 1996; Moran et al., 1993). Prolonged genome erosion may occasionally lead to the loss of symbionts and their replacement by new bacterial lineages (McCutcheon et al., 2019), resulting in repeated cycles of symbiont acquisition, loss, and replacement (Bennett & Moran, 2015; Perreau & Moran, 2022). In contrast, facultative symbionts are not essential for host survival but can provide benefits such as protection or nutritional complementation (Hedges et al., 2008; Teixeira et al., 2008). They retain features of free-living bacteria and are often environmentally acquired or horizontally transmitted (Russell et al., 2003; Sandström et al., 2001) and may serve as an evolutionary reservoir for new obligate symbioses.

In contrast to this canonical pattern, kissing bugs (Reduviidae: Triatominae) represent a notable exception among obligate blood-feeding insects. Over 150 kissing bug species are believed to rely on extracellular gut-associated bacteria acquired via coprophagy rather than intracellular, vertically transmitted mutualists (Beard et al., 2002; Eichler & Schaub, 2002; Orantes et al., 2018). Their microbiomes vary across species, populations, and developmental stages, though they typically converge on a dominant taxon in adults (Brown et al., 2020; Teal et al., 2023). Historically, the actinobacterium *Rhodococcus rhodnii* was considered the sole indispensable nutritional symbiont of triatomines (Eichler & Schaub, 2002). However, recent studies indicate more complex associations, particularly in Rhodniini, including *Enterococcus faecalis*, *Kocuria* sp., and even known intracellular symbionts from the genus *Wolbachia* and *Symbiopectobacterium* (Eberhard et al., 2022; Filée et al., 2022.). The genus *Symbiopectobacterium* stands out as an emerging symbiotic clade with broad ecological and evolutionary flexibility. This genus, closely related to the plant-pathogenic *Pectobacterium*, includes bacteria capable of transitioning between plant-associated niches and a variety of animal hosts, including insects and nematodes (Martinson et al., 2020). Some strains can exist extracellularly and be cultured outside the host. The type species, *Symbiopectobacterium purcellii*, described from the leafhopper *Empoasca decipiens* (Nadal-Jimenez et al., 2022), and the distinct S-MEX strain from another *Empoasca* species (Gunasekaran et al., 2026), likely represent early-stage facultative symbionts. At the opposite end of the spectrum, the *Symbiopectobacterium* symbiont of the bulrush bug *Chilacis typhae* is a vertically transmitted obligate intracellular mutualist housed in specialized bacteriocytes (Kuechler et al., 2011), consistent with long-term coevolution and functional integration. This ecological diversity is mirrored in genome size variation, ranging from large genomes (∼4–5 Mb) typical of facultative bacteria to reduced genomes (∼1.5 Mb).

The presence of *Wolbachia* and *Symbiopectobacterium* in some *Rhodnius* species is reminiscent of the system described in another group of obligate haematophagous heteropterans, the bed bugs (Cimicidae) (Hypša et al., 2025). Among 13 tested species, *Symbiopectobacterium* co-occurred with *Wolbachia* in two species and was the sole symbiont in three others. These recent findings bring us back to questions about the complexity of symbiosis in Rhodniini, as well as the origins and functions of their symbionts, particularly *Symbiopectobacterium* and *Wolbachia*. Specifically, the extent of *Symbiopectobacterium* association across the genus *Rhodnius,* as well as its nature and localization in hosts, remains unclear. Nutritional symbionts in obligate haematophages are typically characterized by an intracellular lifestyle, whereas Triatominae have long been regarded as relying primarily on extracellular symbiosis. Despite decades of investigation, with *Rhodnius prolixus* serving as a key model for insect physiology and anatomy (Nunes□da□Fonseca et al., 2017; Ons, 2017), there is no evidence for a stable intracellular nutritional symbiont corresponding to the canonical system observed in other obligate blood-feeding insects, namely a vertically transmitted symbiont housed in bacteriomes or specialized regions of the gut. The repeated detection of *Symbiopectobacterium* in *R. prolixus* raises the question of whether such a system exists and has remained undetected, or whether *Symbiopectobacterium* maintains an alternative symbiotic strategy in Triatominae.

A different set of questions arises for *Wolbachia* associated with Rhodniini. These bacteria are widespread and abundant across insects, living almost exclusively intracellularly, either as parasites or obligate mutualists. Such intimate associations can lead to the transfer of bacterial genes or even large genomic regions into the host genome, a phenomenon that has also been suggested for *Rhodnius*-associated *Wolbachia* (Eberhard et al., 2022). Finally, as observed in other insect symbioses, *Wolbachia* and *Symbiopectobacterium* have been shown to coordinate their metabolic activities in Cimicidae. Reports of these two genera in *Rhodnius* species raise the possibility of similar coexistence and metabolic complementarity, further underscoring the complexity of the microbiome in these hosts.

In this study, we address these questions to provide new insights into the complexity and function of symbiotic associations in the genus *Rhodnius*. We test the hypothesis that *Symbiopectobacterium* associated with triatomines represents an early stage of symbiotic integration, possibly reflecting a transition from an extracellular to an intracellular lifestyle. We use *Rhodnius prolixus*-associated *Symbiopectobacterium* as a model to characterize its distribution, genomic diversity, and evolutionary relationships across the genus *Rhodnius*, and to evaluate how its genomic features may reflect emerging functional roles. We also examine its localization within host tissues to determine whether it establishes residence within host cells, a key step toward obligate symbiosis. Additionally, we assess the co-occurrence and genomic content of other symbionts, including *Wolbachia*, *Arsenophonus*, and *Rhodococcus*, to identify potential patterns of metabolic complementation. By addressing these questions, we aim to gain a deeper understanding of the unique symbioses of Triatominae and their potential implications for the vectorial capacity of *Trypanosoma cruzi*, the causative agent of Chagas disease.

## Material and methods

### *Symbiopectobacterium* genome GA colony

Scaffolds from the *Rhodnius prolixus* genome assembly (BioProject PRJNA1138944), generated from an adult male originating from a laboratory colony, were screened for bacterial sequences. The details on DNA extraction, HiFi and Hi-C library construction, sequencing, data processing and genome assembly performed using Vertebrate Genome Project (VGP) workflows (Lariviere et al., 2026; Rhie et al., 2021) are described in detail in Habib et al. (2026). Potential bacterial scaffolds in the obtained *R. prolixus* assembly were identified by the BLASTn algorithm (Altschul et al., 1990), querying 10-kb per scaffold against the NCBI nucleotide database. Two scaffolds (1,368,827 bp and 2,665,613 bp) with BLASTn hits to Genus *Symbiopectobacterium* were selected for downstream analysis. Attempts to close the *Symbiopectobacterium* genome using Hi-C and HiFi read mapping and comparison with the *Symbiopectobacterium purcellii* reference genome were unsuccessful, likely due to repetitive phage elements flanking the two scaffolds. The draft genome has been deposited in GenBank (accession GCF_050472005).

### Screening and assembly of *Rhodnius* metagenomic data

SRA data for 36 *Rhodnius* samples (project accession PRJNA429761; Supplementary Table 1) were downloaded and assembled using SPAdes v3.15.3. A BLAST database was then created for each assembly using the BLAST+ suite v2.16.0+ (Camacho et al., 2009). To screen the assemblies for the presence of focal symbionts, we generated query datasets from the genomes of *Arsenophonus triatominarum* (GCF_001640365.1) and *Rhodococcus rhodnii* (GCF_008011915.1) by extracting all annotated coding sequences (CDSs). For *Symbiopectobacterium*, we aimed to maximize representation of the genus by downloading all publicly available genomes from the NCBI database and extracting all unique CDSs, yielding a dataset of 76,514 sequences. The three query datasets were screened against each assembly using BLASTN, retaining up to five hits per query sequence. Candidate symbiont contigs were identified based on these BLAST results. At this stage, any contig containing at least one hit to a symbiont query was considered a potential symbiont-derived contig. To validate the taxonomic origin of these candidate contigs, we performed a reciprocal ("back-BLAST") analysis. All CDSs predicted from the candidate contigs were compared against the NCBI nt database. Only contigs that returned best matches to the corresponding focal symbiont were retained for subsequent analyses. BUSCO v5.8.2 (Simão et al., 2015) was used to assess the completeness of the draft genomes.

### Phylogenetic analyses

To determine the phylogenetic placement of *Symbiopectobacterium*, an initial 16S rRNA gene-based analysis was conducted, followed by phylogenomic analysis. For the 16S rRNA gene analysis, sequences from multiple hosts obtained from previous studies were compiled and aligned using Clustal Omega v1.2.2 (Sievers & Higgins, 2018) in Geneious Prime® 2023.0.1 (Kearse et al., 2012). Phylogenetic reconstruction was conducted using PhyML v3.0 (Guindon et al., 2005) with the Maximum Likelihood method and 100 bootstrap replicates. The best-fit substitution model was determined using the Smart Model Selection (SMS) algorithm implemented in the PhyML online platform, applying the Bayesian Information Criterion.

For the phylogenomics, single-copy orthologs were identified using OrthoFinder v2.4.0 (Emms & Kelly, 2019). The orthologous sequences were aligned with Clustal Omega v1.2.2 and concatenated into a combined dataset. The resulting alignment was manually inspected in Geneious Prime® 2023.0.1 (Kearse et al., 2012), and all positions containg any gaps were automatically removed before phylogenetic inference. Bayesian phylogenetic reconstruction was performed using PhyloBayes (Lartillot et al., 2013), using the CAT-Poisson model with a discrete gamma distribution of rate variation across sites (4 categories), with posterior probabilities used as measures of branch support. The analysis was conducted on an amino acid alignment comprising 26 taxa and 41,864 sites. Convergence between two independent MCMC chains was assessed using the PhyloBayes utilities *tracecomp* and *bpcomp*. Topological convergence was evaluated using bipartition discrepancies (*bpcomp*; maxdiff = 0.16, meandiff = 0.021). Final phylogenetic trees were visualized using iTOL (Letunic & Bork, 2021). In addition to *Symbiopectobacterium*, phylogenetic analyses were conducted for *Arsenophonus*, *Wolbachia*, and *Rhodococcus*. These analyses followed the same workflow as the genome-based *Symbiopectobacterium* tree, but the supplementary phylogenies, included primarily to provide taxonomic context, were inferred with the faster RAxML approach.

### Genome annotation and reconstruction of metabolic pathways

Metabolic functions were evaluated using the Kyoto Encyclopedia of Genes and Genomes (KEGG) database (Kanehisa, Sato, Kawashima, et al., 2016). KEGG ortholog identifiers (K numbers), linking genes to metabolic pathways, were assigned to all coding sequences (CDSs) annotated by PROKKA using the BlastKOALA server (Kanehisa, Sato, & Morishima, 2016). These annotations were then mapped onto KEGG-defined biosynthetic pathways, with a focus on B vitamin synthesis and secretion systems, playing important role in host-symbiont interactions. Given the relatively low coverage of most of the draft genomes, genes missing from the vitamin biosynthetic pathways were manually searched for in the raw sequencing data to ensure their absence was not due to incomplete assembly. Annotation of CRISPR arrays and cas genes was performed with the CRISPRCasFinder program using the default parameters (Couvin et al., 2018).

### Localization of *Symbiopectobacterium* in *Rhodnius prolixus*

Individuals used for this part of the study originate from a laboratory population previously confirmed to be positive for *Symbiopectobacterium* through PCR screening and amplicon sequencing. To determine whether *Symbiopectobacterium* resides intracellularly, guts were dissected from 20 *R. prolixus* individuals and fixed in 4% paraformaldehyde solution for 24 hours. To reduce tissue autofluorescence, the guts were subsequently put in 6% hydrogen peroxide for 10 days. After the quenching, the guts were dehydrated by an ethanol-chloroform series and embedded in paraffin. Serial sections (6 µm thick) were prepared using a rotary microtome and mounted on silicon-coated slides, which were stored in a dry environment until fluorescence *in situ* hybridization was performed. Two probes specific for *Symbiopectobacterium* were designed using Geneious Prime ® 2023.0.1 (Kearse et al., 2012) to target the 16S rRNA gene (FITC-AAGGGCACAACCTCCAAA, and FITC-CTTCTGCGAGTCACGTCAATCAG). Hybridization was conducted following the protocol described by Nováková et al. (2015). We used general EUB338 probe as a positive control and the antisense probe of EUB338, as a negative control for nonspecific binding (Amann et al., 1990; Wallner et al., 1993). Fluorescent signals were visualized using an Olympus FV3000 laser scanning confocal microscope (Olympus Corporation, Tokyo, Japan). Images were captured at 200x, 400x, and 600x magnification and processed using FV31-SW software.

Gut ultrastructure was examined by transmission electron microscopy (TEM) following high-pressure freezing, freeze substitution, resin embedding, and ultrathin sectioning of dissected adult guts (n = 12). To assess the presence of *Symbiopectobacterium* in the hemolymph, DNA extracted from hemolymph and corresponding whole-body samples of fifth-instar nymphs and adults was screened by. Detailed protocols are provided in Supplementary Methods.

## Results and Discussion

### Symbiopectobacterium *genome*

Two *Symbiopectobacterium* contigs, totaling 4,034,440 bp with a GC content of 50.3%, were recovered from the *Rhodnius prolixus* metagenomic assembly (BioProject PRJNA1138944). They likely represent a nearly complete bacterial genome, although it remains unclosed due to the presence of repetitive phage regions at the end of the scaffolds. BUSCO analysis assessed its completeness at 95% using the *Enterobacterales_odb12* dataset. The assembly, with 80.5% coding density, contains 5,050 predicted coding sequences (CDSs) and reveals a highly dynamic genome structure, including 793 transposases, 709 phage-related elements, 21 ribosomal genes, and over 3,048 predicted pseudogenes.

This apparent genomic plasticity is further supported by an extensive repertoire of DNA repair mechanisms and homologous recombination pathways, as well as the presence of a Tad secretion system. Together with core genes encoding the Type III secretion system (T3SS) and the flagellar export machinery, these features suggest a relatively recent establishment of this symbiont, hereafter referred to as *Symbiopectobacterium* RP GCF_050472005. This interpretation is consistent with the general characteristics of other *Symbiopectobacterium* strains as bacteria exhibiting multiple independent transitions from plant pathogen–related ancestors to associations with animal hosts (Gunasekaran et al., 2026; Kuechler et al., 2011; Martinson et al., 2020; Nadal-Jimenez et al., 2022). The genus is widely distributed among arthropods, with roles ranging from facultative to obligate symbiosis, often involving nutrient provisioning (Hypša et al., 2025; Martinson et al., 2020).

#### Metabolic capacity

As a bacterium with a genome size comparable to that of its free-living relatives, *Symbiopectobacterium* RP GCF_050472005 retains much of its ancestral metabolic capacity, including complete biosynthetic pathways for riboflavin (vitamin B2), pyridoxal (B6), pantothenate (B5), biotin (B7), and a nearly complete pathway (E3.1.3.1 missing) for folate (B9) (Fig. 1 A and Supplementary Table 2). The intermediate stage of evolution toward a host-adapted symbiont is further supported by its secretion system repertoire. The genome encodes only core components of the Type III secretion system (T3SS), whereas it lost Type IV (T4SS), often associated with horizontal gene transfer and host manipulation, and Type VI (T6SS) secretion systems mediating interbacterial competition in environmentally exposed bacteria (Unni et al., 2022). In the stable and protected environment of host tissues, these systems become unnecessary and are frequently eliminated during genome reduction. Their absence suggests reduced reliance on environmental competition and horizontal exchange.

**Figure 1.**
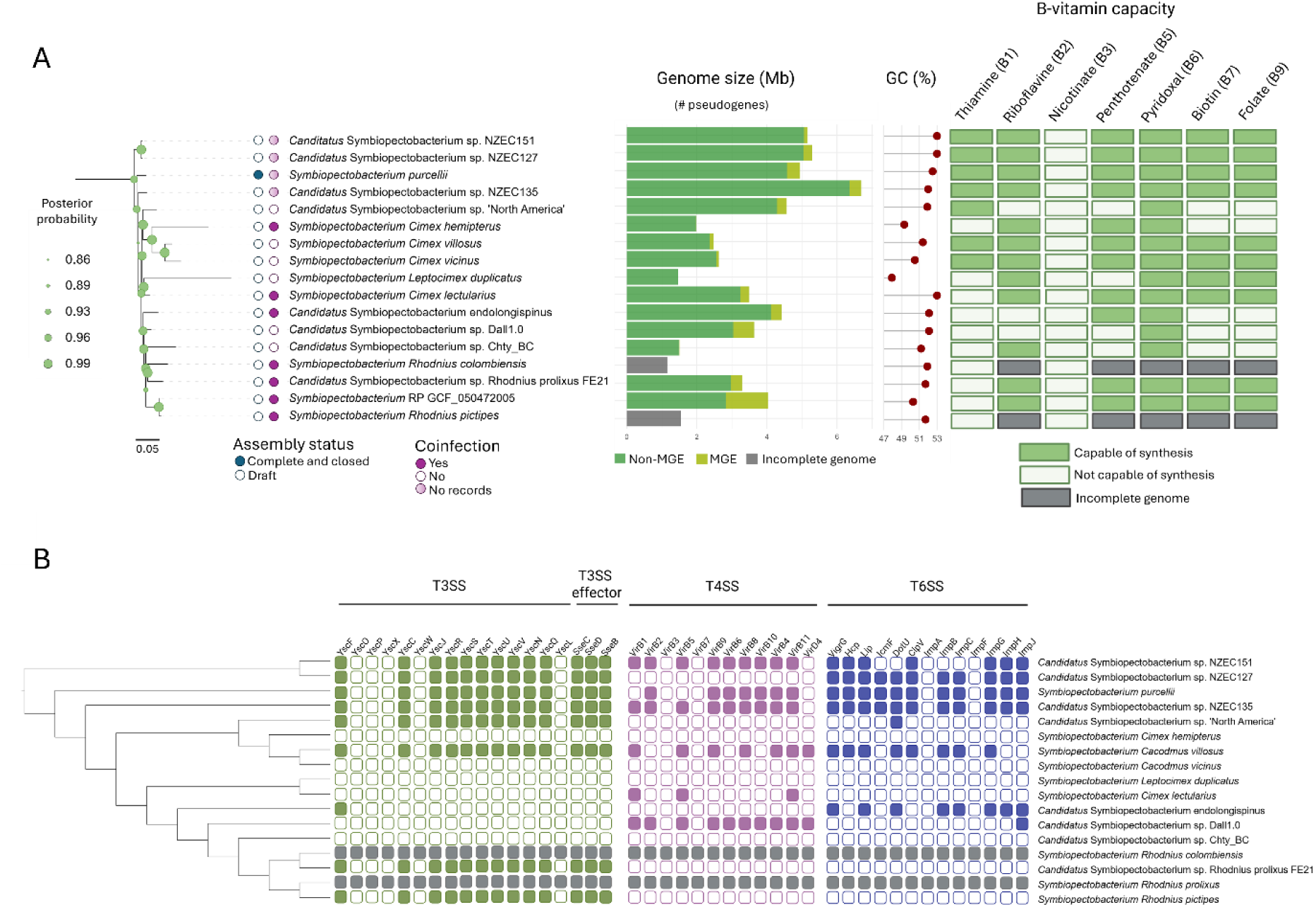
Comparative genomics of *Symbiopectobacterium.* (A) Genome-based phylogeny (full phylogeny including outgroups shown in Supplementary Fig. 1) with the *Symbiopectobacterium* clade expanded; posterior probabilities are indicated. Assembly status and co-infection are denoted by colored circles (see legend). Horizontal bars show genome size (Mb), with the fraction of mobile genetic elements (MGEs; phages and plasmids) in yellow; numbers in brackets indicate pseudogene counts. GC content (%) is shown as a lollipop plot. The heatmap displays predicted B-vitamin biosynthetic capacity (BlastKOALA), with filled boxes indicating presence. B: Core components of T3SS (green), T4SS (magenta), and T6SS (blue) as predicted by BlastKOALA are shown. Absence of genes is indicated by an empty square, presence with a colored square. The relationship of *Symbiopectobacterium* is shown with a cladogram based on the core genome phylogeny in (A).

The secretion repertoire of the *Symbiopectobacterium* RP GCF_050472005 contrasts with the *Symbiopectobacterium* strains confirmed as obligate endosymbionts, such as the strain from *C. typhae* and several cimicid-associated strains, which have lost all major secretion systems (Fig. 1B; Hypša et al., 2025; Kuechler et al., 2011). The T3SS is typically associated with host cell invasion and early stages of intracellular establishment. Recently emerged endosymbionts often retain this machinery to facilitate interactions with host tissues (Dale et al., 2001). Once stable intracellular residency is established, particularly within bacteriomes, the T3SS becomes dispensable and is subsequently lost (Siozios et al., 2024). The retention of T3SS in the *Symbiopectobacterium* RP GCF_050472005, therefore, suggests ongoing host interaction and incomplete symbiotic integration.

Consistent with this interpretation is the absence of a complete CRISPR–Cas system. The loss of CRISPR-based defense is commonly observed in vertically transmitted symbionts and may facilitate the accumulation of mobile genetic elements during transitional stages (Burstein et al., 2016; Jiang et al., 2013; Zaayman & Wheatley, 2022). The high content of mobile elements in *Symbiopectobacterium* RP GCF_050472005 thus likely reflects active genome remodeling during host adaptation.

#### Tissue localization

Localization within host tissues is a key indicator of the nature of host-symbiont interactions. Symbionts with nutritional roles are typically intracellular, residing in bacteriomes or associated with specific regions of the host gut epithelium. In *Symbiopectobacterium*, intracellular localization has been conclusively demonstrated or indicated only in a limited number of cases (Kuechler et al., 2011; Martinson et al., 2020), whereas for many insect-associated lineages the spatial organization of the symbiosis remains unresolved.

To determine the localization of *Symbiopectobacterium* in *Rhodnius prolixus* and to assess its potential intracellular lifestyle, we performed fluorescence in situ hybridization (FISH), transmission electron microscopy (TEM), and hemolymph screening. FISH analysis detected *Symbiopectobacterium* RP GCF_050472005 exclusively in the posterior midgut (Fig. 2C-D), where signals were observed within gut epithelial cells. No bacteria were detected in the rectal ampulla or anterior midgut (Fig. 2A-B). This localization is notable, as the posterior midgut is the primary site of digestion and nutrient absorption in triatomines (Garcia & Azambuja, 1991; Kollien & Schaub, 2000). The restriction of *Symbiopectobacterium* to this region, together with its capacity for B-vitamin biosynthesis, is therefore consistent with a putative nutritional role.

**Figure 2.**
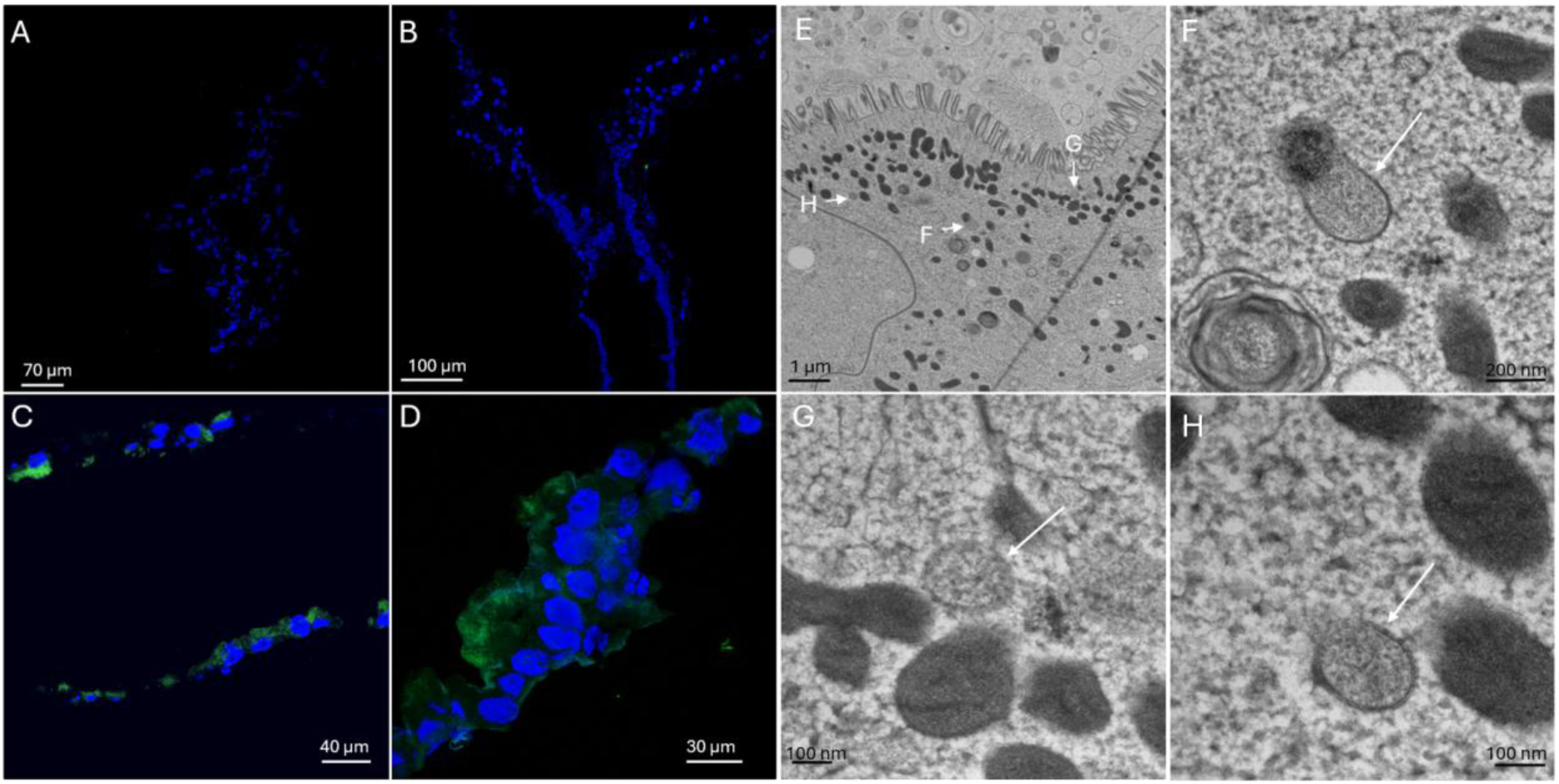
Confocal and transmission electron microscopy (TEM) of gut sections of *Rhodnius prolixus*. Fluorescence *in situ* hybridization (FISH) using a *Symbiopectobacterium* RP GCF_050472005–specific probe shows bacterial localization. Hybridization signals for DAPI (cyan) label eukaryotic nuclei, while the specific probe labeled with FITC (green) marks bacterial cells. (A) rectal ampulla; (B) transition between anterior and posterior midgut; (C, D) posterior midgut sections. (E) TEM image of posterior midgut tissue showing intracellular bacterial cells; arrows indicate *Symbiopectobacterium* RP GCF_050472005. (F-H) higher-magnification TEM views of bacterial cells. (E) arrows indicate *Symbiopectobacterium* cells; (F-H) higher-magnification views of bacterial cells.

However, the distribution of this bacterium varied strongly among the samples. In some individuals, strong fluorescent signals were concentrated near the luminal (apical) side of epithelial cells (Fig. 2C-D), whereas in others, these signals were sparse or unevenly distributed along the gut epithelium (Supplementary Fig. 2). TEM observations confirmed the intracellular presence of *Symbiopectobacterium* but did not reveal the dense bacterial accumulations observed by FISH (Fig. 2E). Only a limited number of rod-shaped bacterial cells, approximately 200 nm in diameter, were detected, primarily near the microvilli inside the posterior midgut epithelial cells. The absence of large intracellular bacterial groups in the TEM samples may reflect differences in feeding status, as these individuals were starved prior to processing, whereas specimens showing higher bacterial loads by FISH had been fed.

The detection of *Symbiopectobacterium* in the hemolymph of several individuals (Supplementary Fig. 3) indicates that the bacteria may be capable of crossing the gut epithelial barrier and entering the hemocoel. This translocation was not consistently observed, suggesting that systemic spread is either variable or transient. Similar observations have been reported in the bean bug *Riptortus pedestris*, where facultative gut-associated bacteria can cross the gut epithelium, enter the hemocoel, and trigger systemic immune responses, including activation of the Toll and IMD pathways and upregulation of antimicrobial peptides (Jang et al., 2024). Although epithelial barrier crossing is often associated with pathogenicity, in *Riptortus pedestris* it did not negatively affect host fitness and instead enhanced resistance to subsequent infections by pathogenic bacteria. The heterogeneous patterns observed here are consistent with a similarly dynamic interaction, rather than a stable obligate endosymbiosis. Occasional systemic exposure may therefore contribute to host immune priming.

These findings indicate that the intracellular lifestyle of *Symbiopectobacterium* in *R. prolixus* is not fixed. Although the bacteria can associate with epithelial cells and may cross the gut barrier, colonization is inconsistent, and no bacteriome-like structures or specialized crypts were observed, suggesting that the association has not reached the level of integration characteristic of obligate endosymbiosis. Preliminary microbiome inheritance experiments (Moons et al., in prep.) indicate that embryos are sterile, arguing against transovarial transmission. Together, these observations suggest that *Symbiopectobacterium* represents a facultative symbiont capable of transient intracellular residence and epithelial translocation, consistent with a transitional stage in the evolution toward stable endosymbiosis.

### Screening of SRA: Symbiopectobacterium and other bacterial associates of the genus *Rhodnius*

*Symbiopectobacterium* has been previously detected in the microbiome of *Rhodnius prolixus* as an incomplete metagenome-assembled genome (MAG) (Eberhard et al., 2022), suggesting that its association with kissing bugs is not incidental. Together with our identification of *Symbiopectobacterium* RP GCF_050472005, which exhibits traits consistent with a transition toward intracellular nutritional symbiosis, this points to a more complex symbiotic landscape in *Rhodnius* species than previously recognized, traditionally centered on *Rhodococcus rhodnii*. These observations raise the question of how widespread *Symbiopectobacterium* is across the genus *Rhodnius* and to what extent it contributes to the diversity of symbiotic associations in these haematophagous insects. To address this, we reassembled 36 publicly available Sequence Read Archive (SRA) datasets representing 17 *Rhodnius* species and performed a comprehensive analysis of their bacterial communities.

We detected *Symbiopectobacterium* sequences in metassemblies of *R. pallescens*, *R. pictipes*, *R. neglectus*, *R. nasutus*, *R. colombiensis*, and *R. prolixus*, indicating a broad distribution of this symbiont within the genus (Fig. 3A). While short contigs of *Symbiopectobacterium* and other bacteria are found across the genus (Supplementary Table 1), the quality of SRA data only allowed for the assembly of two *Symbiopectobacterium* MAGs. Compared to the genome of *Symbiopectobacterium* RP GCF_050472005, the MAGs are more fragmented and are approximately 2.5 times smaller, 1.4Mb and 1.7Mb. Their GC content is 52.3%, and genome completeness, as assessed with BUSCO using the *pectobacteriaceae_odb12* dataset, ranged from 33% to 77% (Fig. 2B). The lower completeness likely reflects the predominance of environmental bacteria in the *pectobacteriaceae_odb12* database. When using the *Enterobacterales_odb12* dataset, these numbers increase, from 43% to 95%. The broad completeness range, along with assembly fragmentation, aligns with the differences found in repetitive content of the three draft genomes. *Symbiopectobacterium* RP GCF_050472005 contains 19 phage regions and 793 transposases. Given the incompleteness of the other *Symbiopectobacterium* draft genomes, comparisons of pseudogene and mobile element content could not be interpreted.

**Figure 3:**
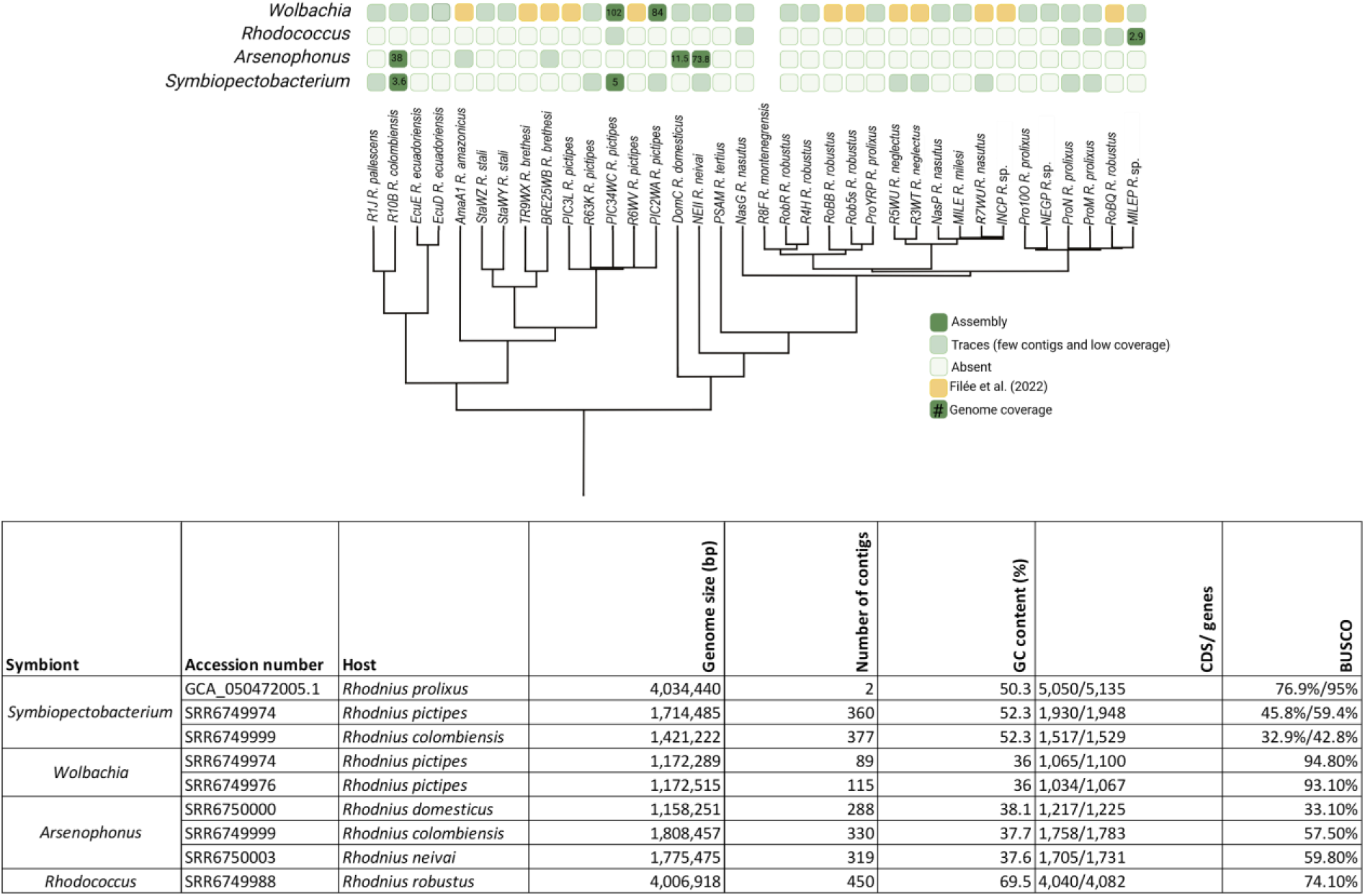
(A) Distribution of the symbionts found throughout the *Rhodnius* genus. Dark green are the draft genomes assembled based on the SRA data from Filée et al. (n.d.). Yellow designates *Wolbachia* draft genomes assembled previously by Filée et al. (n.d.). Embedded numbers stand for an average coverage. (B) Basic characteristics *of* the assembled MAGs. BUSCO analysis was done based on the closest taxonomic group available. Phylogeny was traced from Filée et al. (2022) figure 1.

In addition to *Symbiopectobacterium*, our analysis recovered three other symbiont-associated taxa, *Wolbachia*, *Arsenophonus*, and *Rhodococcus*. Among these, *Wolbachia* was the most prevalent, with sequences detected across all analyzed *Rhodnius* datasets (Fig. 3A), consistent with previous findings by Filée et al. (2022). Two of the *Wolbachia* assemblies recovered here, both from *R. pictipes*, are comparable in structure and genome statistics to those reported by Filée et al. (2022). Phylogenetic analyses based on 35 orthologous genes among 46 taxa (Supplementary table 3) place *Rhodnius*-associated *Wolbachia* in a well-supported monophyletic clade within supergroup F, closely related to the strains from the bedbug *Cimex lectularius* (wCle) and the filarial nematode *Madathamugadia hiepei* (Supplementary Fig. 4). This phylogenetic proximity to wCle may reflect relatively recent horizontal transfer events between distantly related hosts, as suggested by Filée et al. (2022). However, the widespread presence of *Wolbachia*-derived sequences across the genus, together with a large integrated region identified in the *R. prolixus* genome (acc. no. GCA_049639745.1) spanning approximately 176 kb, points to repeated past infections and a long-term association. This interpretation is consistent with evidence for recurrent *Wolbachia*-to-*Rhodnius* gene transfers reported by Filée et al. (2022) and Mesquita et al. (Mesquita et al., 2015), supporting a scenario in which ancient and ongoing interactions have shaped the symbiotic landscape of the genus.

The second bacterium, the genus *Arsenophonus*, was detected in five species, *R. colombiensis*, *R. domesticus*, *R. neivai*, *R. neglectus*, and *R. nasutus*, and the third genus, *Rhodococcus*, occurred in *R. nasutus* and *R. robustus* (Fig. 3A). Both these genera are well-known associates of triatomines. *Arsenophonus triatominarum* was discovered in *Triatoma infestans* and is reported mainly from *Triatoma* species (da Mota et al., 2012; Hypša & Dale, 1997). *Rhodococcus rhodnii* has long been considered the main symbiont of *Rhodnius* species (Ben-Yakir, 1987; Brecher & Wigglesworth, 1944; Gilliland et al., 2022), whereas *R. triatominae* has been associated with the genus *Triatoma* (Yassin, 2005). All the *Arsenophonus* MAGs described in this study clustered with *A. triatominarum*. (Supplementary Fig. 5). Likewise, the *Rhodococcus* MAG obtained from *R. robustus* showed closest affinity to *R. rhodnii* and grouped within a larger clade that also includes *R. triatominae*, described from another *Triatoma* species (Supplementary Fig. 6). Remarkably, despite all the datasets belonging to *Rhodnius* species, *Rhodococcus*-related contigs were recovered from only four SRA datasets, which was unexpected given its presumably established role as a core symbiont in this genus. However, PCR screening conducted by Filée et al. (2022) detected *Rhodococcus* in all analyzed samples, supporting the view that this symbiont remains widespread in *Rhodnius* despite its limited recovery from the sequencing datasets examined here. The low recovery of *Rhodococcus* sequences may be influenced by the difficulty of lysing and extracting DNA from members of this genus (Nahar et al., 2021).

#### Origins and putative roles

We analyzed the phylogenetic position of the *Symbiopectobacterium* RP GCF_050472005 together with the other *Symbiopectobacterium* sequences assembled from the other *Rhodnius* SRA data, and the strain reported by Eberhard et al. (2022). The proteome matrix comprised 198 single-copy orthologues, totaling 68,600 aligned amino acid residues.

All *Rhodnius*-associated strains formed a well-supported monophyletic group, with *Symbiopectobacterium* infecting *C. typhae* (*Candidatus* Symbiopectobacterium sp. Chty_BC) being the closest relative (Fig. 1A). The strains associated with *R. colombiensis* and *R. prolixus* reported by Eberhard et al. (2022) clustered together and were both placed on a longer branch relative to other *Rhodnius*-infecting strains. Notably, the two strains infecting *R. prolixus* individuals did not form a monophyletic cluster, suggesting independent acquisition events. However, this pattern could also result from methodological or taxonomic factors. In addition to potential artefacts associated with metagenome-assembled genomes (MAGs), such as incomplete assemblies or hybrid sequences, species identification within *Rhodnius* can be challenging due to morphological similarity among taxa and the possible presence of cryptic species or hybrids. The 16S rRNA gene phylogeny (Supplementary Fig. 7) further supports the placement of *Symbiopectobacterium* RP GCF_050472005 detected in the *R. prolixus* colony within the Triatominae-associated *Symbiopectobacterium* clade. It forms a monophyletic clade with sequences obtained from other *R. prolixus* individuals and *Dipetalogaster maximus*. As noted previously by da Mota et al. (2012), *Symbiopectobacterium* is not restricted to a single host genus and likely infects a wide range of Triatominae species.

Genome characteristics vary markedly among *Symbiopectobacterium* strains. While GC content remains relatively stable (48–53%), genome size, pseudogene content, and the abundance of mobile elements differ substantially (Fig. 1A), reflecting divergent ecological strategies. A clear distinction emerges between plant-associated strains and host-restricted endosymbionts. For example, the strain infecting the bulrush bug *C. typhae* is a confirmed obligate symbiont (Kuechler et al., 2011), characterized by a highly reduced genome (1.4 Mb) and placement on a long phylogenetic branch, consistent with long-term host dependence. Similarly reduced genomes and even longer branches are observed in strains associated with the bedbugs *Cimex hemipterus* and *Leptocimex duplicatus*. In contrast, plant-associated strains isolated from potato tubers retain much larger genomes (5–6.7 Mb), indicative of a less intimate, likely environmental lifestyle.

Within *Rhodnius*, *Symbiopectobacterium* strains show varying degrees of host adaptation. The relatively short branch lengths of strains associated with *R. prolixus* and *R. pictipes* suggest more recent associations, whereas the strain from *R. colombiensis* occupies a longer branch, indicating a more ancient relationship. This is further supported by its lower GC content, comparable to or even below that of cimicid-associated symbionts, a reduced number of pseudogenes, and a more advanced deterioration of the T3SS (Fig. 1B), all consistent with a more progressed stage of symbiotic integration. Taken together, these patterns align with a trajectory of transition from environmental bacteria to obligate endosymbionts, in which early stages are marked by the proliferation of mobile elements and pseudogenization, followed by progressive genome reduction and stabilization in long-term intracellular associations. The diversity observed among *Rhodnius*-associated strains thus likely reflects ongoing and potentially independent transitions toward more intimate symbiosis.

When assessing the possible role of *Symbiopectobacterium* in Triatominae, comparisons with other blood-feeding hemipterans are informative. In bed bugs (family Cimicidae), *Symbiopectobacterium* has been reported in several species (Hypša et al., 2025). While in some hosts it co-occurs with the obligate symbiont *Wolbachia*, making its role less clear, in others it is the sole detected symbiont, strongly suggesting a function as an obligate nutritional mutualist. This parallel between Triatominae and Cimicidae becomes more apparent when considering *Wolbachia*: in both groups, it is the most prevalent symbiont, whereas *Symbiopectobacterium* either co-occurs with it or, in some cases, appears to complement or replace its nutritional role. This possibility is particularly relevant in *R. pictipes* and *R. colombiensis*, where *Wolbachia* lacks the capacity for biotin biosynthesis (Filée et al. 2022). Although the incompleteness of the *Symbiopectobacterium* genomes recovered here precludes a robust assessment of biotin biosynthetic capacity, previously characterized *Symbiopectobacterium* symbionts from *Rhodnius* retain this pathway. Therefore, it remains plausible that the *Symbiopectobacterium* strains detected in these species contribute to biotin provisioning, potentially compensating for the loss of this function in *Wolbachia*, although this cannot be confirmed with the current genome assemblies. A comparable pattern is observed in the *R. prolixus* colony from which we obtained the insects for this study, in which individuals harbor both *Rhodococcus* and *Symbiopectobacterium* RP GCF_050472005. This Symbiopectobacterium retains a complete biotin synthesis pathway, whereas this pathway is incomplete in *R. rhodnii* (Gilliland et al., 2023). Together, these cases support a direct nutritional role for *Symbiopectobacterium*, at least in systems where metabolic complementarity is evident. More broadly, in Triatominae, this role is further supported by its intracellular localization within the gut epithelium and signs of early genome degradation, whereas in Cimicidae, it is reinforced by its occurrence as the sole symbiont in some host species.

## Conclusion

Our results reveal that *Symbiopectobacterium* is a widespread and evolutionary dynamic associate of *Rhodnius* species, displaying genomic and ecological features consistent with ongoing transition toward intracellular nutritional symbiosis. The combination of genome plasticity, mobile element expansion, pseudogenization, and partial secretion system loss alongside retained B-vitamin biosynthesis suggests that these bacteria are undergoing progressive adaptation to host association while still maintaining characteristics of facultative lifestyles.

At the same time, the variable intracellular localization of *Symbiopectobacterium* within posterior midgut epithelial cells, its occasional detection in the hemolymph, the absence of specialized bacteriomes, and its inconsistent occurrence across *Rhodnius* species indicate that this association has not reached the degree of host dependence and integration characteristic of obligate endosymbioses. Comparative analysis further suggests that *Rhodnius*-associated strains occupy different positions along the symbiotic continuum, ranging from recently acquired facultative associates to more host-restricted lineages exhibiting signatures of advanced genome reduction.

The recurrent co-occurrence of *Symbiopectobacterium* with *Wolbachia*, *Arsenophonus*, and *Rhodococcus* also supports the existence of a flexible and potentially metabolically complementary microbial network rather than dependence on a single obligatory partner. Altogether, these findings challenge the traditional view of Triatominae as relying exclusively on extracellular symbiosis and establish *Symbiopectobacterium* as a valuable model for studying the evolutionary emergence of obligate intracellular mutualism in blood-feeding insects.

## Acknowledgment

This research has been funded by the Czech Science Fundation (grant numbers 21-10185M to EN and to 24-10943S VH). We acknowledge the BC CAS core facility LEM, supported by the Czech-BioImaging large RI project (LM2023050 and OP VVV CZ.02.1.01/0.0/0.0/18_046/0016045 funded by MEYS CR), for their support with obtaining scientific data presented in this paper.

## Data availability

The metagenome-assembled genomes (MAGs) generated in this study have been deposited in GenBank under BioProject accession **PRJNA1480505**.

