## Supplementary Figures for "Caught in transition: facultative intracellularity and genome evolution of *Symbiopectobacterium* in *Rhodnius* species"

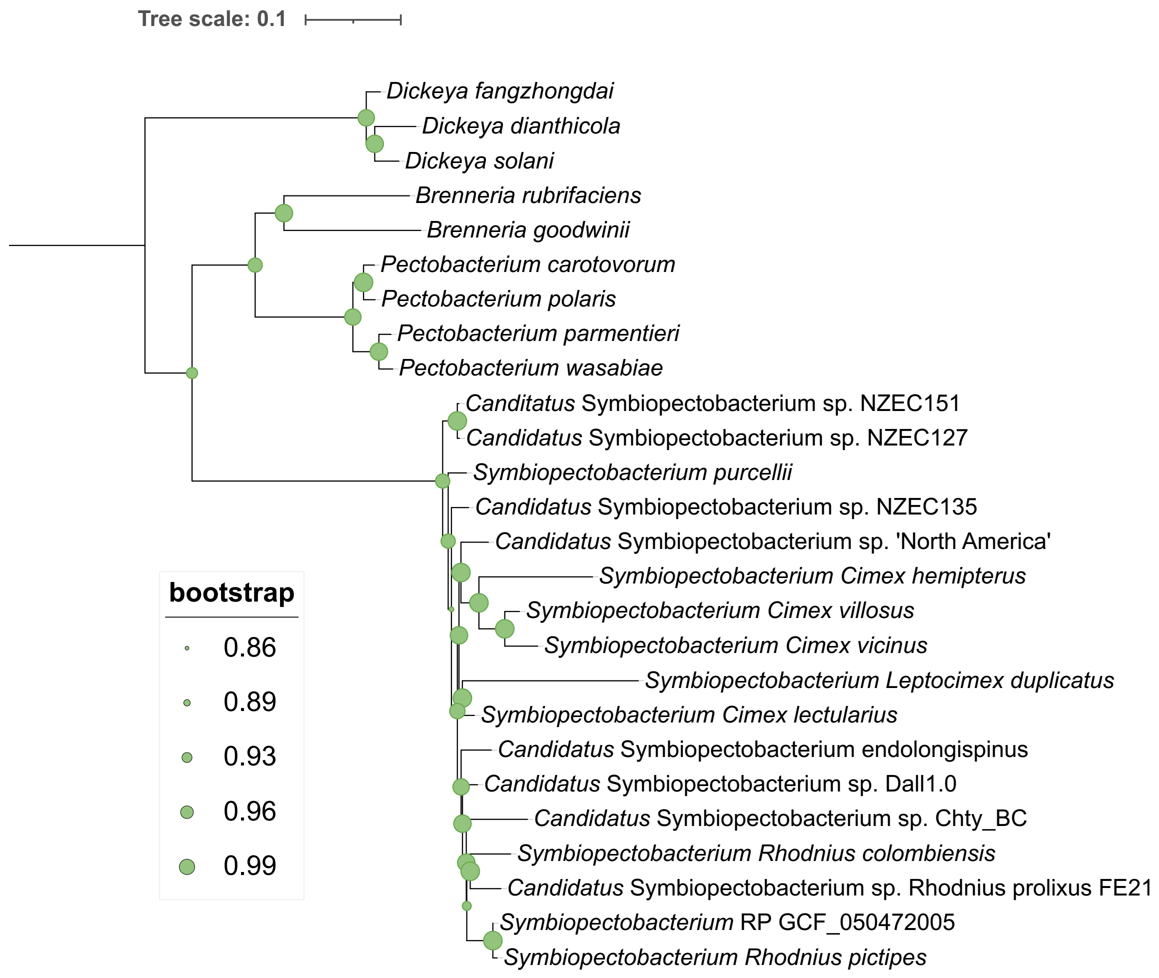


**Supplementary Figure 1. Bayesian genome-based phylogeny of *Symbiopectobacterium* sp.** conducted on an amino acid alignment comprising 26 taxa and 41,864 sites.

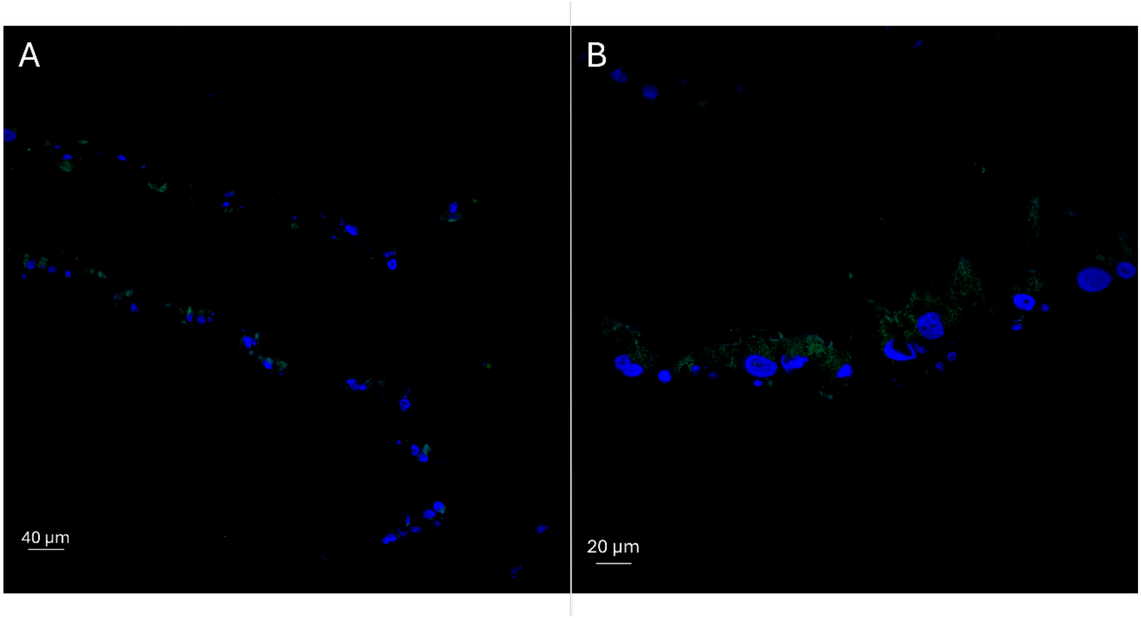


**Supplementary Figure 2. Confocal microscopy of gut sections of *Rhodnius prolixus*.** Fluorescence in situ hybridization (FISH) using a *Symbiopectobacterium* RP GCF_050472005–specific probe shows bacterial localization. Hybridization signals for DAPI (cyan) label eukaryotic nuclei, while the specific probe labeled with FITC (green) marks bacterial cells.

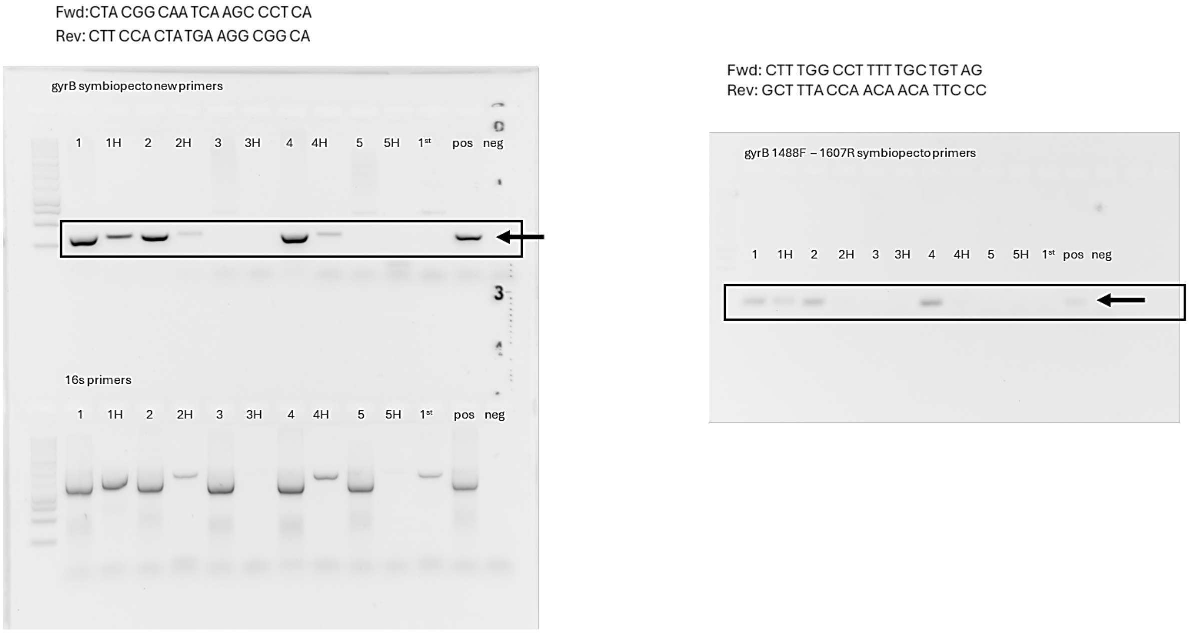


**Supplementary Figure 3. Detection of *Symbiopectobacterium* DNA in hemolymph samples by PCR.** PCR amplification of hemolymph-derived bacterial DNA using *gyrB*-specific primers (upper and right panel) and 16S rRNA primers (lower panel). Amplicons were visualized by agarose gel electrophoresis. Positive and negative controls are shown, and arrows indicate expected PCR products.

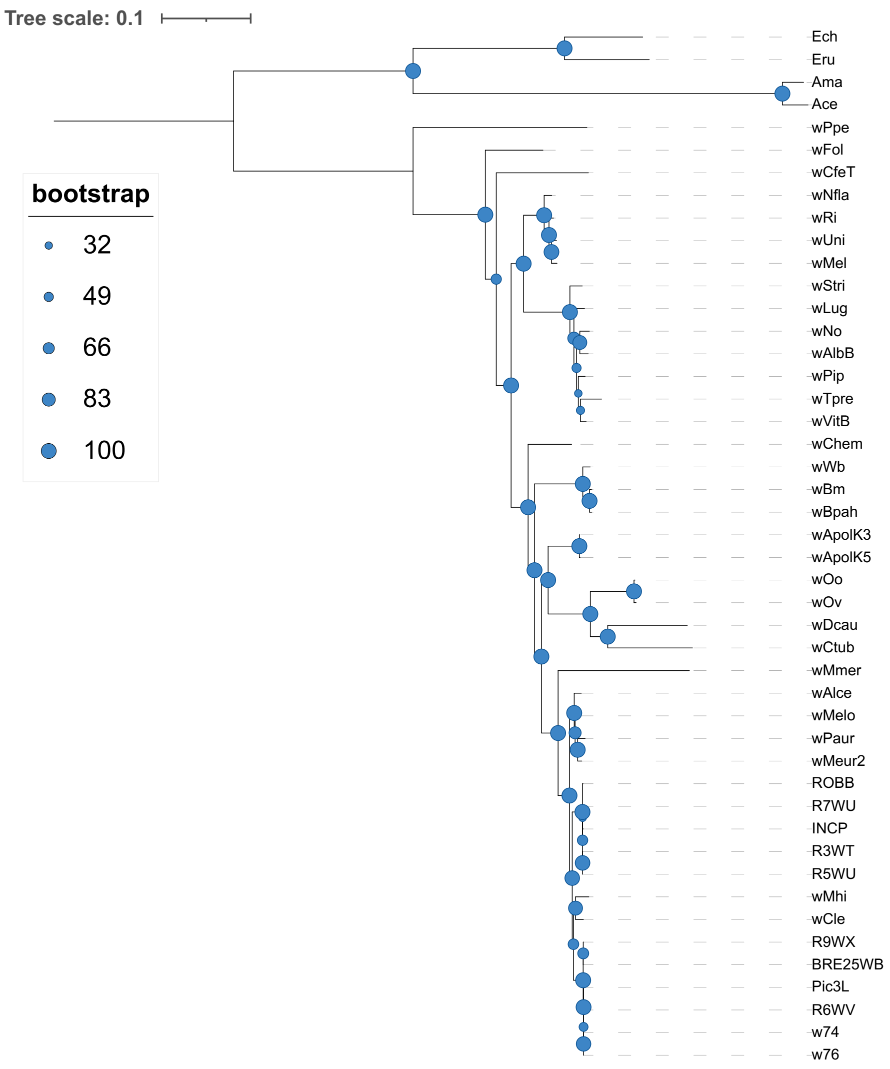


**Supplementary Figure 4. ML genome-based phylogeny of *Wolbachia* species.**
A maximum-likelihood phylogenetic tree was inferred from a concatenated alignment of 35 single-copy orthologs identified by OrthoFinder, comprising 10,659 amino acid sites across *Wolbachia* genomes and selected reference taxa. Phylogenetic reconstruction was performed using RAxML with the DAYHOFF amino acid substitution matrix and the GAMMA model of rate heterogeneity. Branch support was assessed using 100 rapid bootstrap replicates. Node labels indicate bootstrap support values, and branch lengths are proportional to the estimated number of substitutions per site. Details on host taxa and accession numbers can be found in Supplementary table 3.

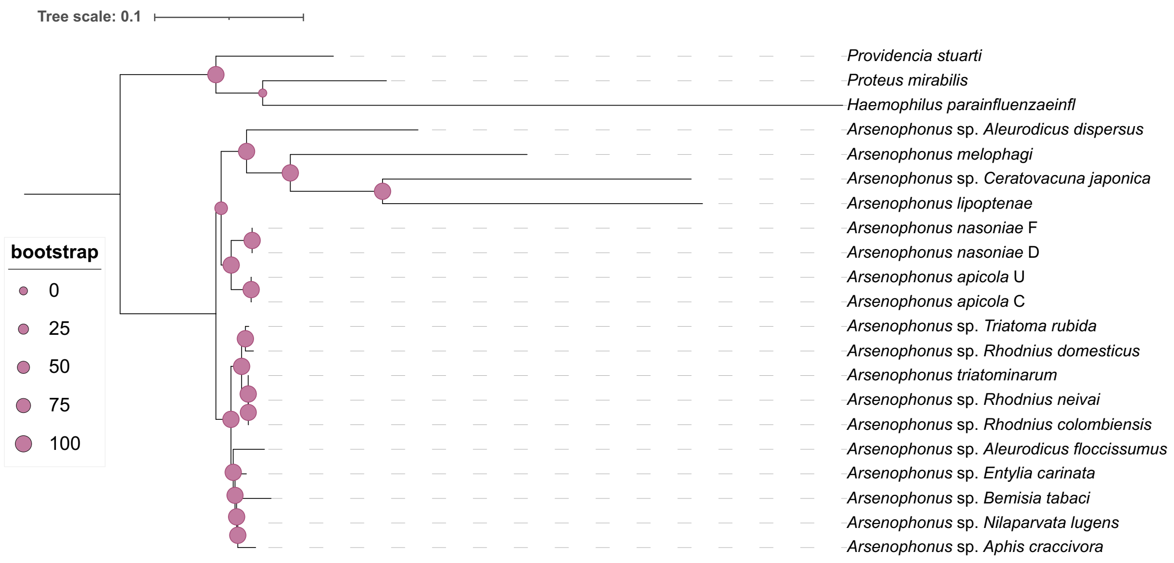


**Supplementary Figure 5. ML genome-based phylogeny of *Arsenophonus* species.**
A maximum-likelihood phylogenetic tree was inferred from a concatenated alignment of 101single-copy orthologs identified by OrthoFinder, comprising 32,506 amino acid sites across *Arsenophonus* genomes and selected reference taxa. Phylogenetic reconstruction was performed using RAxML with the DAYHOFF amino acid substitution matrix and the GAMMA model of rate heterogeneity. Branch support was assessed using 100 rapid bootstrap replicates. Node labels indicate bootstrap support values, and branch lengths are proportional to the estimated number of substitutions per site.

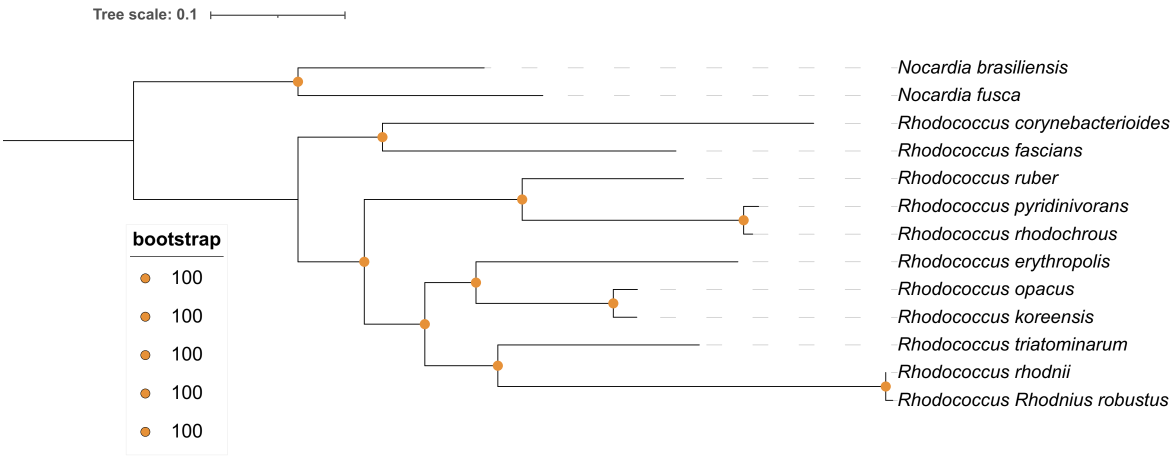


**Supplementary Figure 6. ML genome-based phylogeny of *Rhodococcus* species.**
A maximum-likelihood phylogenetic tree was inferred from a concatenated alignment of 182 single-copy orthologs identified by OrthoFinder, comprising 49,757 amino acid sites across *Rhodococcus* genomes and selected reference taxa. Phylogenetic reconstruction was performed using RAxML with the DAYHOFF amino acid substitution matrix and the GAMMA model of rate heterogeneity. Branch support was assessed using 100 rapid bootstrap replicates. Node labels indicate bootstrap support values, and branch lengths are proportional to the estimated number of substitutions per site.

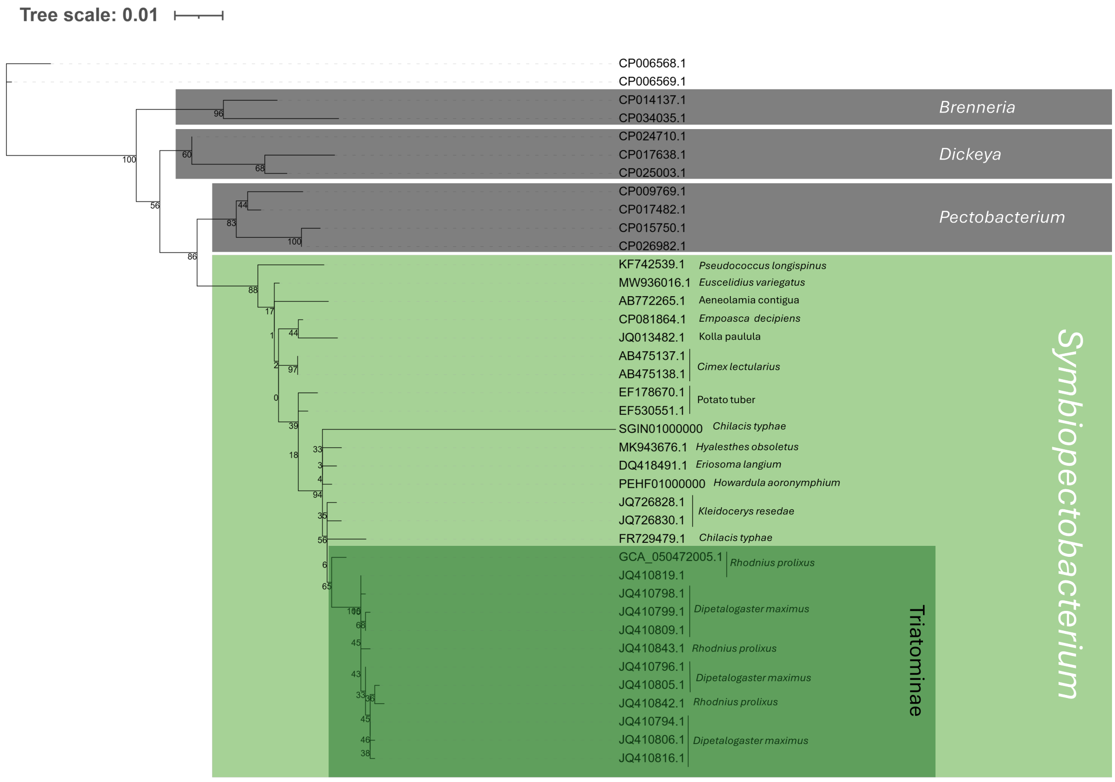


**Supplementary Figure 7. ML phylogeny of *Symbiopectobacterium* based on 16S rRNA gene sequences.**

Phylogenetic tree constructed using the maximum-likelihood method under the HKY85 model with 1055 aligned nucleotide sites. Bootstrap values from 100 replicates are shown at branch nodes. The scale bar indicates the number of substitutions per nucleotide site.
