## Supplementary Methods for "Caught in transition: facultative intracellularity and genome evolution of *Symbiopectobacterium* in *Rhodnius* species"

**Transmission electron microscopy (TEM)**

The entire gut, including the crop and rectal ampule, was dissected in PBS from 12 adult individuals. The tissue was high-pressure frozen using an EM ICE (Leica, Wetzlar, Germany) high-pressure freezer. Freeze substitution EM AFS2 (Leica, Wetzlar, Germany) was carried out in 2% osmium tetroxide diluted in 100% acetone at -90 °C for 16 h, and then warmed up at a rate of 5 °C per hour to remain at -20 °C for 14 h, and finally warmed up again at the same rate to a final temperature of 4 °C. Samples were rinsed three times in anhydrous acetone at room temperature and infiltrated stepwise in acetone mixed with SPI-pon resin (SPI) (acetone : SPI ratios of 2:1, 1:1 and 1:2, for 1 h at each step). The samples, now in pure resin, were polymerized at 60 °C for 48 h. Sections were prepared using a Ultracut UCT (Leica) microtome and collected on 300/400 mesh copper grids (SPI). Staining was performed using ethanol solution of uranyl acetate for 30 min and in lead citrate for 20 min. Images were obtained using a 1400 Flash (JEOL Ltd., Tokyo, Japan) transmission electron microscope.

***Symbiopectobacterium* screening hemolymph**

Samples were collected from fifth-instar nymphs and adult individuals. Hemolymph was obtained by removing one leg and gently compressing the abdomen to form a droplet, which was collected using a pipette. Approximately 4-7 µl of hemolymph was collected per individual and immediately transferred to 400 µl of ZymoBIOMICS Lysis solution. After hemolymph collection, the corresponding whole individuals were also placed in separate tubes containing ZymoBIOMICS Lysis solution. DNA extraction was performed using the ZymoBIOMICS DNA Miniprep Kit (Zymo Research, Irvine, CA, USA) according to the manufacturer’s instructions. Extracted DNA from both hemolymph and whole-body samples was screened using universal 16S rRNA primers as well as two *Symbiopectobacterium*-specific primer sets targeting the *gyrB* gene (gyrB 1488F: FWD CTTTGGCCTTTTTGCTGTAG, REV GCTTTACCAACAACATTCCC; gyrB new: FWD CTACGGCAATCAAGCCCTCA, REV CTTCCACTATGAAGGCGGCA). PCR products were visualized by gel electrophoresis.
